# Genomics Analysis Illuminates Morphology, Ecology, Phenology and Distribution of Two Cryptic *Atrytonopsis* Skippers (Hesperiidae: Hesperiinae)

**DOI:** 10.64898/2026.06.16.732465

**Authors:** Steven J. Cary, Simon M. Doneski, Jing Zhang, Qian Cong, Nick V. Grishin

## Abstract

The Hesperiine genus *Atrytonopsis* Godman, 1900, occurs broadly across the American Southwest. *Atrytonopsis margarita* (Skinner, 1913) and *Atrytonopsis python* (W. H. Edwards, 1882) have look-alike appearances, concurrent flights, and geographic distributions which converge in New Mexico. Their similar wing markings and intertwined taxonomic history has made it challenging to fully understand the identity and occurrence of each. Burns (2015) revealed differences in genitalia, clarifying that they are distinct species. Genomic DNA analysis of more than 100 specimens now illuminates their genetic uniqueness, phylogenetic relationship, field identification challenges and details of their geographic distributions.

---

*Atrytonopsis python* (W. H. Edwards 1882, henceforth Python) and *Atrytonopsis margarita* (Skinner 1913, henceforth Margarita) have a relationship befitting their phenotypic similarity. Python was initially described from Arizona and had been on the books for almost three decades before Margarita was first recognized in New Mexico and described as a full species (Edwards 1882, Skinner 1913). Then Margarita spent most of the 20th Century demoted to the rank of a form, synonym, subspecies or variety of Python, or of the similar *Atrytonopsis cestus* (W. H. Edwards 1884) (Miller and Brown 1981). Burns (2015) discovered a unique pattern of sclerotization of female genitalia which convincingly distinguished the two organisms, then he restored Margarita to a full species. Male genitalia of Python and Margarita were not deemed distinctive.

After differentiating these two skippers in the laboratory, Burns noted differences in wing characters visible to the naked eye: “Wing spots are white in *A. margarita* instead of light yellow, as they are on both wings dorsally and on the forewing ventrally, in *A. python*.” Burns also contrasted dorsal overscaling of both wings by long hairs that are “warm yellow” on *A. python* but “paler yellow to gray” on *A. margarita*. These differences help distinguish Margarita and Python in the field and in collections, but difficulties remain.

Burns went on to describe the geographic distribution of each skipper based on collection locations for specimens he examined: “southwestern New Mexico and Arizona” for Python and “west Texas and in much of New Mexico” for Margarita. This was major progress in our geographic understanding, but it left details to be refined. For example, do their ranges overlap? Knowing the distribution of each species in better detail might illuminate our understanding of the evolution of species and biogeographic boundaries, while aiding in identification of other specimens, photographs or sightings.

For Margarita and Python, total genome DNA analysis of more than 100 specimens offered opportunities to (1) document species-level phylogenetic distinctions, (2) enhance the ability to distinguish them in the field, and (3) establish more detailed maps of their distributions than have heretofore been available.

## MATERIALS AND METHODS

### Material Studied

Accomplishing our three objectives required examination and determination of dozens of Python and Margarita specimens from throughout their combined inhabited area. Specimens were selected to cover as much geographic area as possible in as much detail as practicable. We obtained specimens for analysis from several institutional collections in the United States: Clyde P. Gillette Museum at Colorado State University (CSU), McGuire Center for Lepidoptera at the University of Florida (MGCL), Los Angeles County Museum (LACM), US National Museum (USNM), University of New Mexico’s Museum of Southwest Biology (MSB), Peabody Museum of Natural History (PMNH), Illinois Natural History Survey (INHS), Carnegie Museum of Natural History (CM), Texas A & M University (TAMU), and Florida Museum of Natural History (UF).

In addition, we sampled three specimens from the personal collection of William Dempwolf, one from the personal collection of Nick V. Grishin, and one from Matthew Brown. Collectively, more than 100 specimens of Margarita and Python were sampled and sequenced. They included the holotype of *A. python* (W. H. Edwards 1882) located at the CMNH, two syntypes of *A. margarita* (Skinner 1913) also at CMNH, plus five “types” of *A. margarita* collected by John Woodgate and located at the PMNH.

### Genomic Analysis

Total genomic DNA was extracted from a leg; genomic libraries were constructed and sequenced using the Illumina next-generation platform according to our published protocol (Li et al. 2010). Exons of protein-coding genes were assembled from the sequence data using our standard computational pipeline (Li et al. 2010) and three phylogenetic trees were constructed using IQ-TREE v1.6.12 with the GTR+GAMMA model (Nguyen et al. 2015): one from the autosomes in the nuclear genome, one based on genes predicted to locate on the Z chromosome, and the third from the mitochondrial genome. For nuclear genome trees, 100,000 codons (300,000 base pairs), which is ∼2% of the total codon dataset, were randomly sampled to reduce computational load. Branch support in the resulting trees was evaluated using 100 replicate samples, each comprising 10,000 randomly selected codons from the total dataset. Phylogenetic trees were constructed for each replicate, and support values (ranging from 0 to 100) correspond to the number of replicates that share the same bipartition as the main tree constructed from 100,000 codons. The mitochondrial genome tree was constructed from all available codons and ultrafast bootstrap support was employed (Hoang et al. 2018). Tree figures were made using FigTree (Rambaut 2018).

### Morphological Analyses

To address field identification challenges, SJC visually examined photos of more than 40 study specimens in relation to their genetic IDs, seeking to confirm Burns’ (2015) naked-eye characters and hoping to identify useful ventral hindwing characters.

## RESULTS

### Genomic Analysis

Results of genomic analyses (Fig. 1) affirm that Python and Margarita are distinct species, as previously determined by Burns (2015) based on female genitalia. First, in the Z-chromosome tree (Fig. 1b), the two species form two prominent clades without a confidently supported tree structure within each clade, and each clade could be approximated by a star tree: nearly equidistant genomes within each clade (representing intra-species variation) with a genetic gap between the clades (representing inter-species variation). The Z chromosome is usually best for species delimitation, and such a pattern is indicative of species-level taxa (Ficarrotta et al. 2022). Second, even in the tree constructed from all protein-coding genes on autosomes, Python and Margarita separate into two clades. The autosome tree gives clues about gene exchange between these species, and the placement of specimens from neighboring populations as sisters to all others indicates that gene exchange correlates with the geographic proximity of these populations, which is expected. The lack of such a correlation in the Z chromosome tree (star clades) indicates more restricted gene exchange on the Z chromosome and argues for a completed speciation process between Python and Margarita. The mitochondrial genome tree shows little genetic differentiation and reveals that mitochondrial haplotypes do not correlate with species (Fig. 1c). This situation is common in butterflies, and therefore mitochondrial DNA, including the COI barcode, should be used with caution in species-delimitation or identification studies.

**Figure 1.**
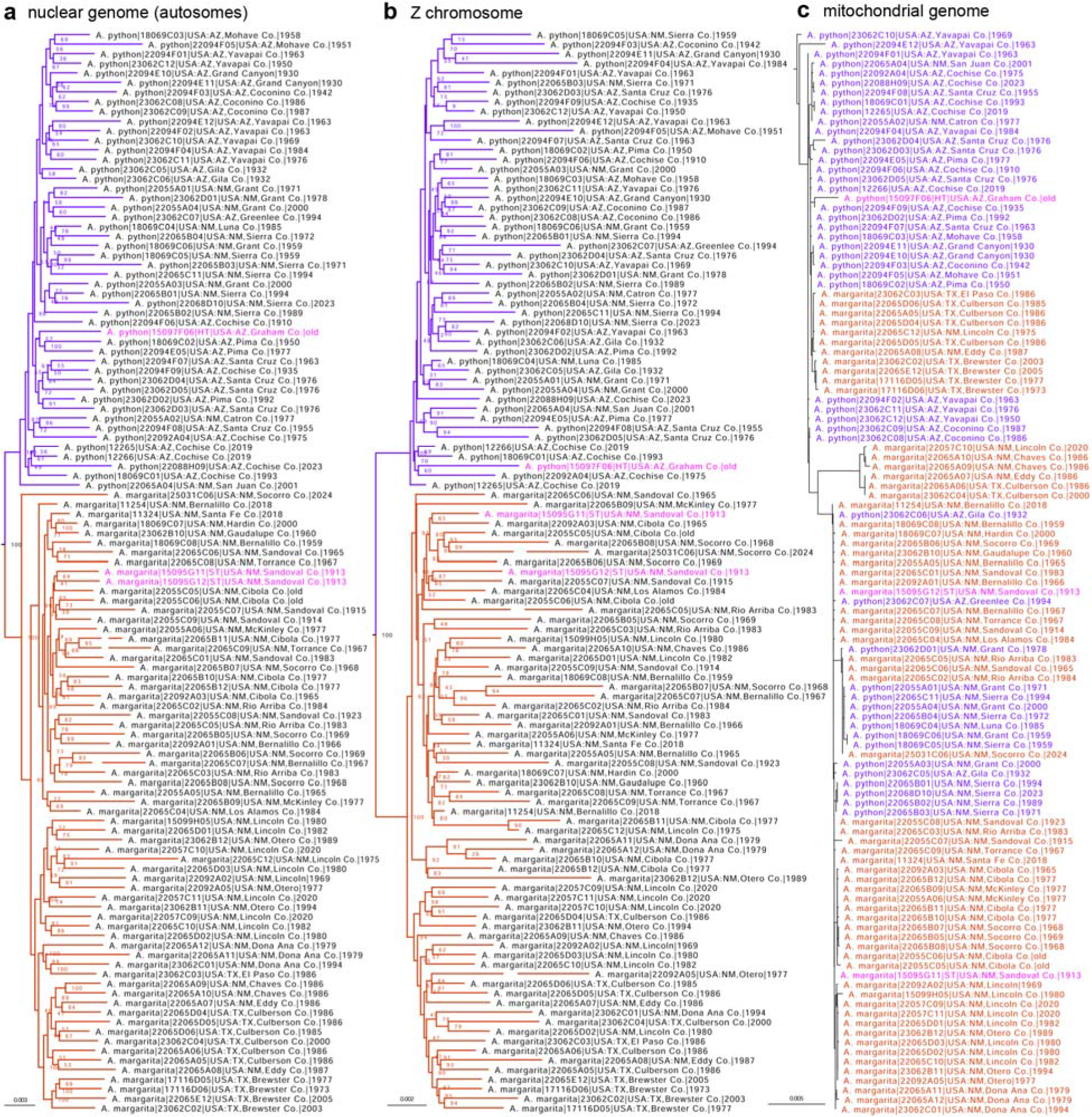
Phylogenetic trees of *A. python* (violet) and *A. margarita* (red) constructed from protein-coding regions in **a)** the nuclear genome (autosomes), based on 925,206 positions, **b)** the Z chromosome, based on 110,019 positions, and **c)** the mitochondrial genome. Name-bearing type specimens are labeled in magenta. Ultrafast bootstrap (Hoang et al. 2018) values are shown at nodes. Gaps in terminal branches indicate that a segment of a branch was cut out to reduce its length (to allow an increase in the font size), i.e., a branch with a gap is longer than shown.

### Morphological Analyses

Typical examples of specimens with Python DNA and Margarita DNA are shown in Figure 2. These medium-sized skippers have a typical forewing length of 17mm and a typical hindwing radius of 13mm. They are variably brown or gray with a standard array of forewing pale spots and a standard array of hindwing maculations which are variably expressed. The two figured specimens convey Python’s warmer, yellower tone compared to Margarita’s generally cool, gray appearance as noted by Burns (2015), but this difference was not consistent among all specimens.

**Figure 2.**
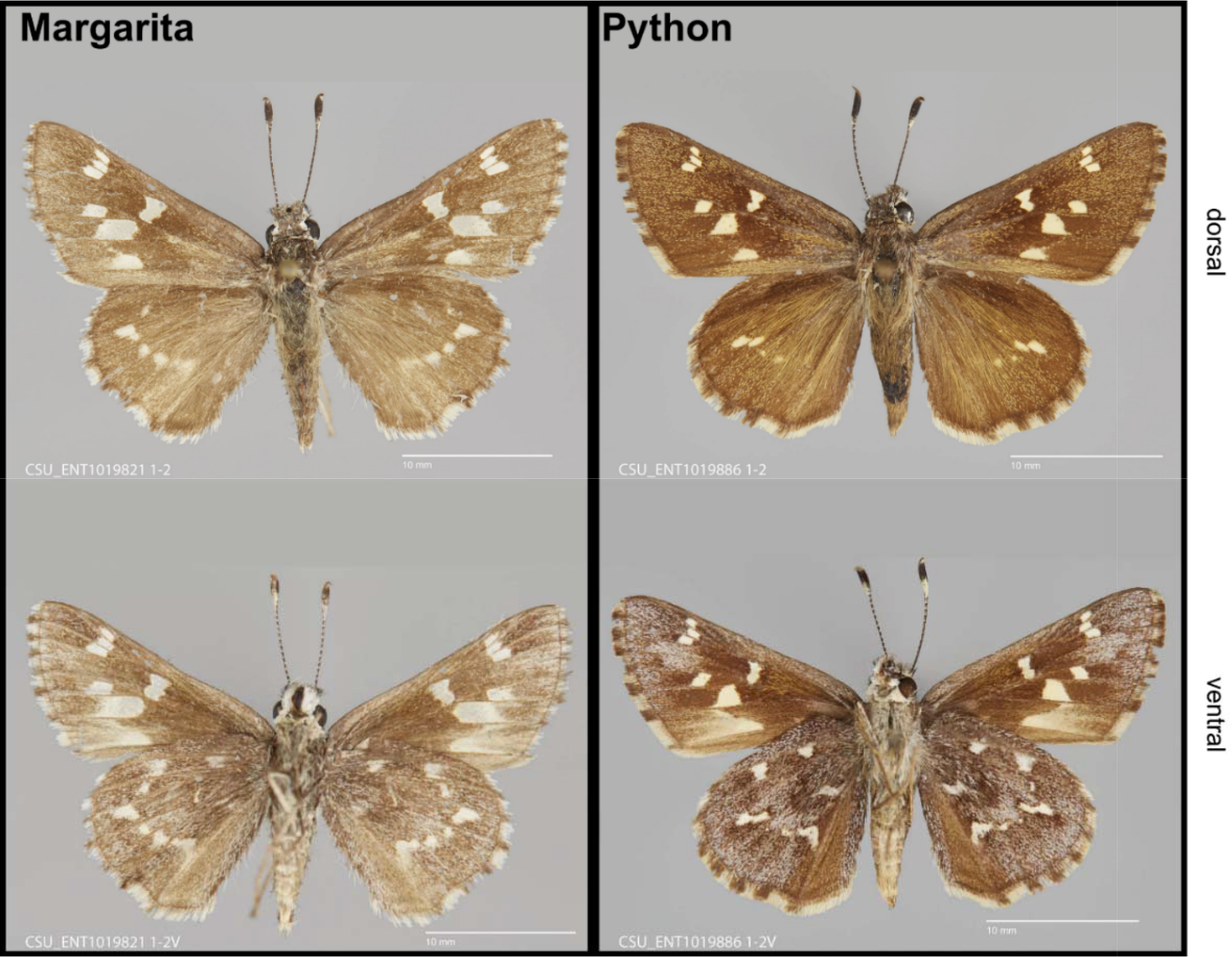
Dorsal and ventral wing surfaces of (A) A. margarita from Jemez Mountains, Rio Arriba Co., NM; May 20, 1983 (coll. R. Holland) and (B) A. python from the Black Range, Sierra Co., NM; May 6, 1989 (coll. R. Holland). Photos courtesy of Chuck Harp.

Prior to DNA analyses, the 40 Holland specimens sampled at CSU were visually examined by SJC, who recorded his tentative naked-eye determination of each specimen as either Python or Margarita. After DNA tests were done, SJC’s naked-eye determinations were compared against genetic identifications. Naked-eye determinations of Margarita matched genetic results only 67 percent of the time (23 of 34). Naked-eye determinations of Python matched genetic results only 50 percent of the time (3 of 6). Several Margarita specimens had yellow-tinted forewing spots and a few Python specimens had untinted forewing spots. Similarly, the overscaling with long hairs was often yellow on Python and paler on Margarita, but not always. Some specimens of both species had long hairs that were dull brown as opposed to yellow or gray.

The ventral hindwing of these two skippers is usually the most colorful and intricately-patterned wing surface. On newly emerged individuals the wash of pale scales can be sky blue, soon fading to gray and white. On study specimens, we found the ventral hindwing’s pale postmedian band to vary in an interesting way. Typically, it is white, but on newly-emerged individuals it can have significant yellow/orange tinting (Fig. 3).

**Figure 3.**
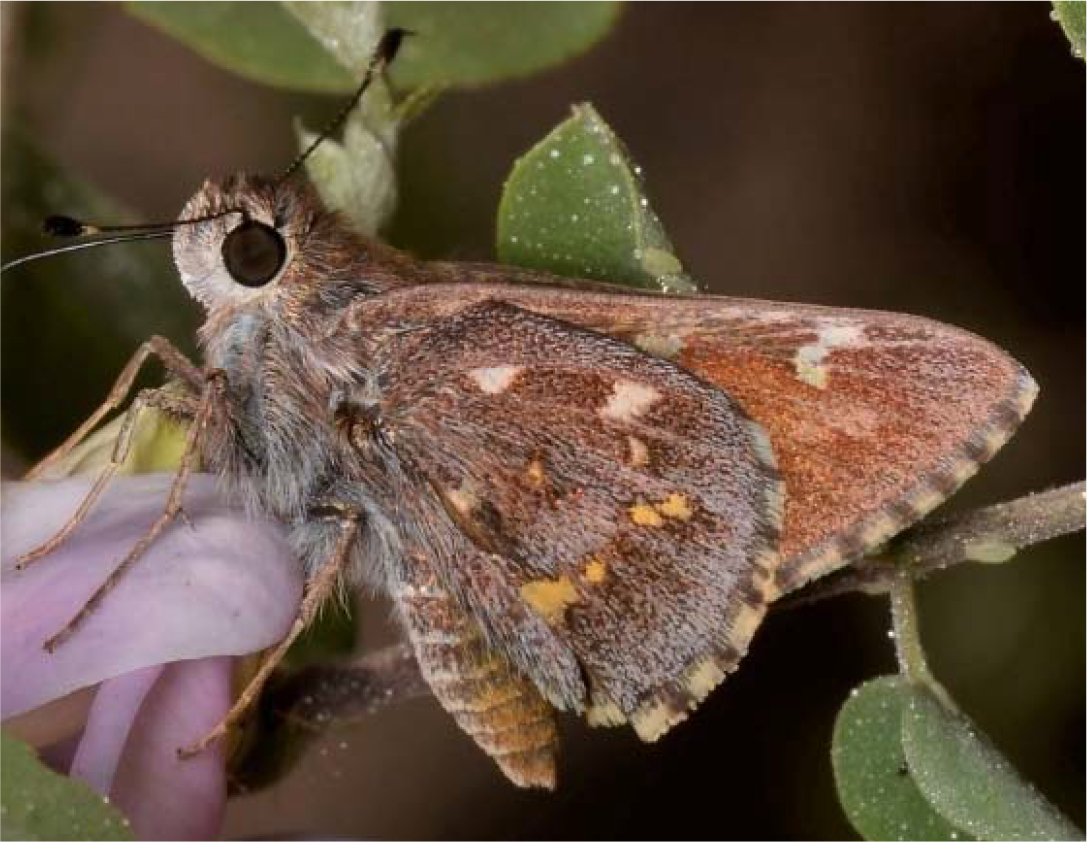
*Atrytonopsis* sp., Water Canyon, Magdalena Mountains, Socorro Co., NM; May 2022 (photo by Matthew Brown).

Examining that character in greater depth on museum specimens of Python and Margarita, as well iNaturalist observations, we found that about ten percent of Margarita specimens and about 20 percent of Python individuals had some yellowing of that postmedian spot band, rendering it not particularly useful in distinguishing the two skippers.

Wing fringes of the two species are also quite striking (Figs. 2, 3). On Python, wing fringes tended to be distinctly marked, with wide black checks against a faint, salmon-colored hue. On Margarita, fringes tended to be dirty white with black checks that were blurred or smeared and often narrow. But there was considerable overlap of these subjective characters among various specimens of the two species, so this character also proved unhelpful.

## DISCUSSION

### Genomics

Genomic work presented above affirms and supplements the work of Burns (2015) by providing an additional way to conclusively distinguish Margarita specimens from Python specimens. This new DNA discriminant then enabled rigorous comparison of known Python against known Margarita regarding wing morphology, use of the landscape, phenology and geography.

### Wing Morphology and Identification Using Naked-Eye Characters

We hoped to enhance the ability to distinguish Python and Margarita in the field or from photographs, but in this regard we were not successful. Our two study subjects are splendid examples of cryptic species: difficult to distinguish from each other without genitalic dissections or DNA analyses.

### Flight Times and Flight Elevations

Of the >100 specimens we analyzed, knowing their genetic identities allowed more precise characterization of Python and Margarita as distinct species in terms of flight periods, altitudinal preferences, and geographic occurrences. For example, within the North American portion of their respective distributions, Python and Margarita adults are essentially synchronic. Charting study specimens with usable dates, Fig. 4 shows that adults of these two skippers fly concurrently in their respective habitats. For each, the one annual flight is equally spread between May and June, with infrequent occurrences in late April and again in July and August.

**Figure 4.**
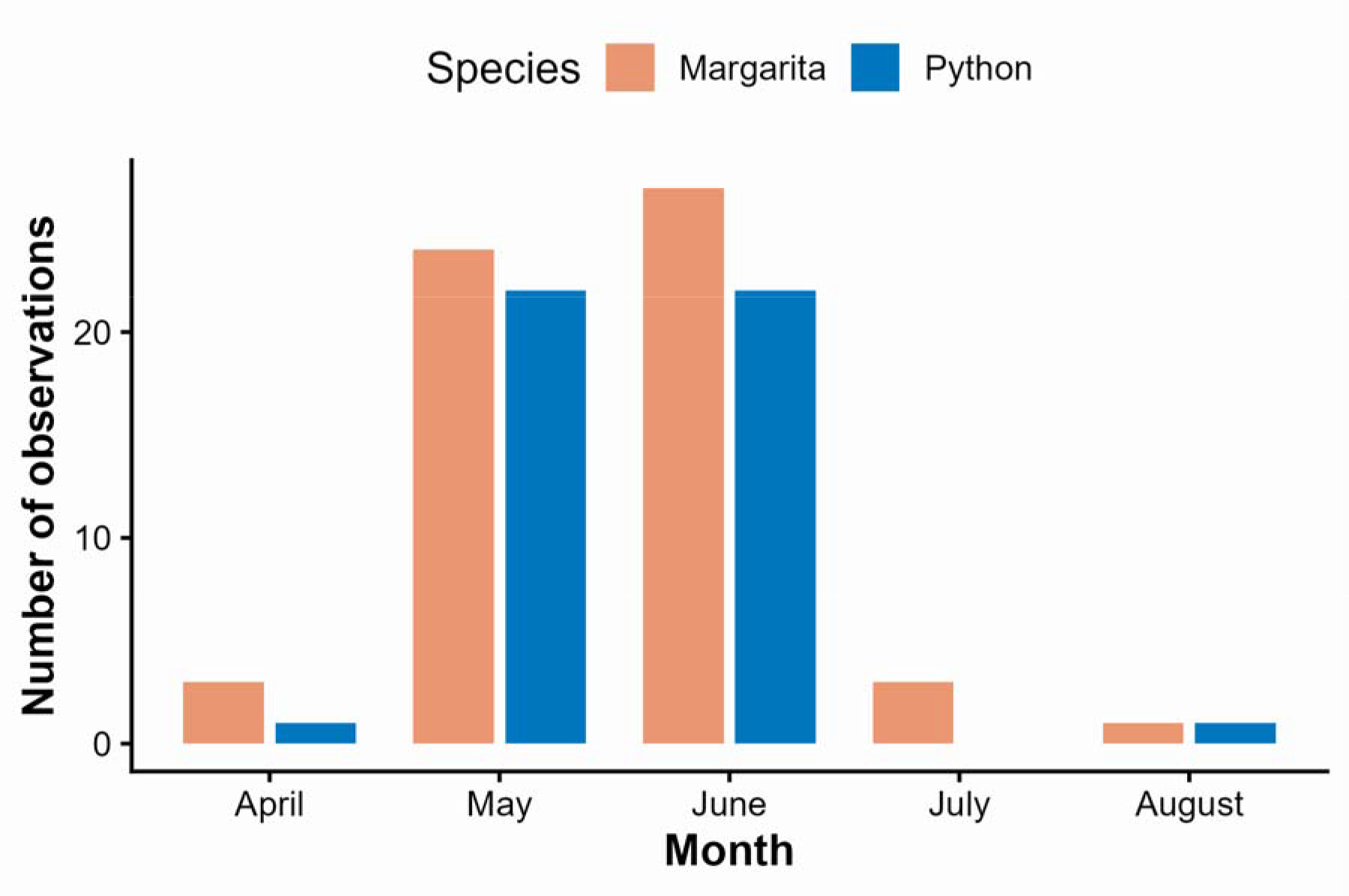
Number of study specimens captured in months of April through August for *A. margarita* (orange, n=58) and *A. python* (blue, n=44).

We also examined topographic elevations at which adults fly. Figure 5 shows numbers of study specimens captured in altitudinal bands spanning <1200m (<4,000ft) to 2900m (9,500ft) elevation. Both species fly primarily between 1200m (4,000ft) and 2450m (8,000ft) elevation, with infrequent lower and higher sightings. For the most part, plant communities at these elevations are dominated by savannas of pines, oaks and grasses (McNally 2020:213). Overall, Margarita occurs across a slightly broader span of altitudes while being concentrated between 1825m (6,000ft) and 2275m (7,500ft). In contrast, Python is almost equally distributed across altitudes between 1200m (4,000ft) and 2275m (7,500ft).

**Figure 5.**
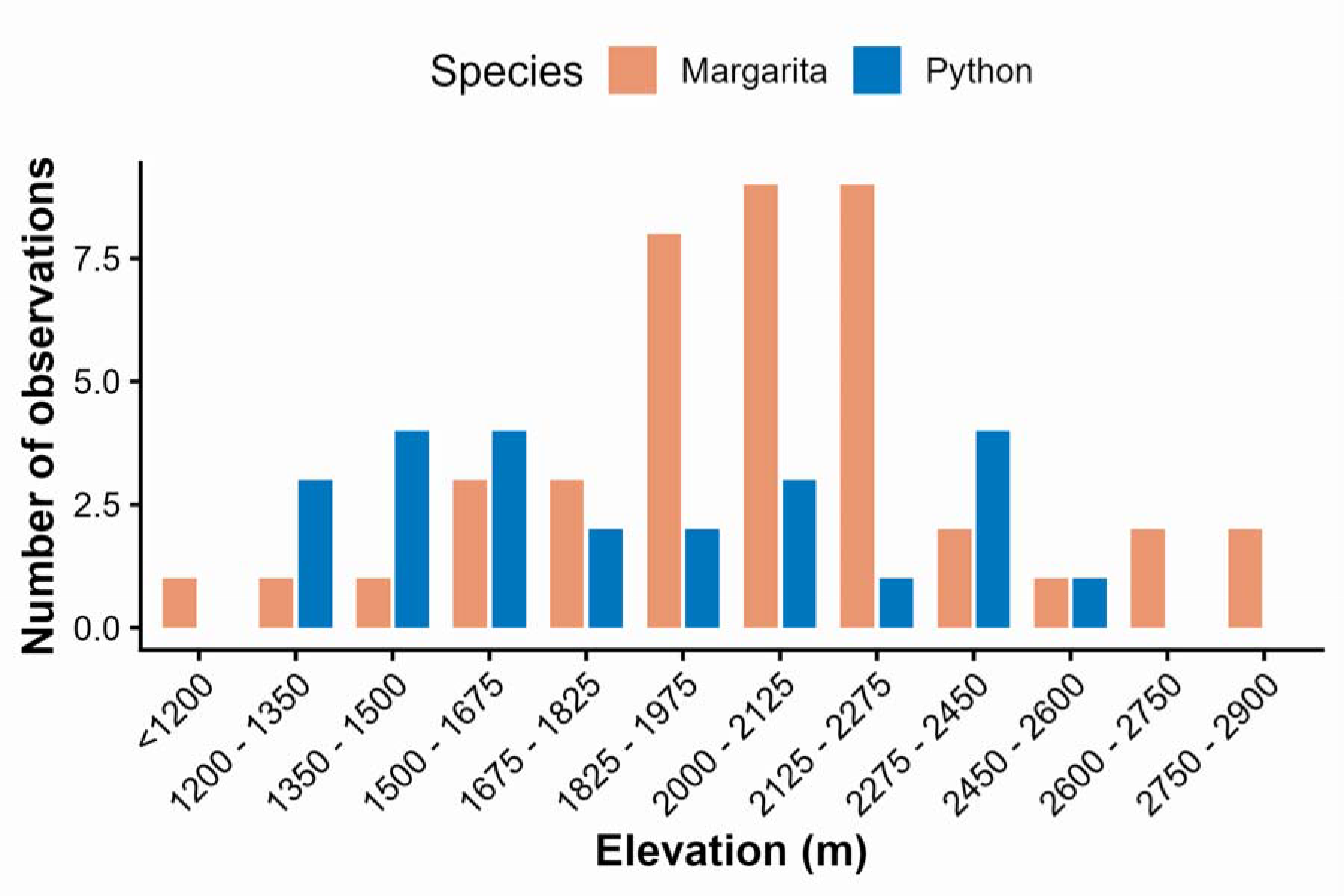
Numbers of study specimens of *A. margarita* (orange, n=42) and *A. python* (blue, n=24) captured at specified elevations.

The two skippers have nearly identical appearances, while flying at the same time at essentially the same altitudes. These similarities help explain why it has been difficult to distinguish them in the field, and why the detailed geographic distribution of each has been difficult to delineate with confidence.

### Geography

Genomic results allow individual occurrences of each species to be mapped with greater precision than was heretofore possible (Fig. 6). Study specimens with Margarita DNA originated in New Mexico and west Texas, but not Arizona; study specimens with Python DNA originated in Arizona and southwest New Mexico, but not Texas. Though very similar in terms of wing morphology, habitat, phenology and ecology, each member of this cryptic pair is most readily distinguished by its unique geography.

**Figure 6.**
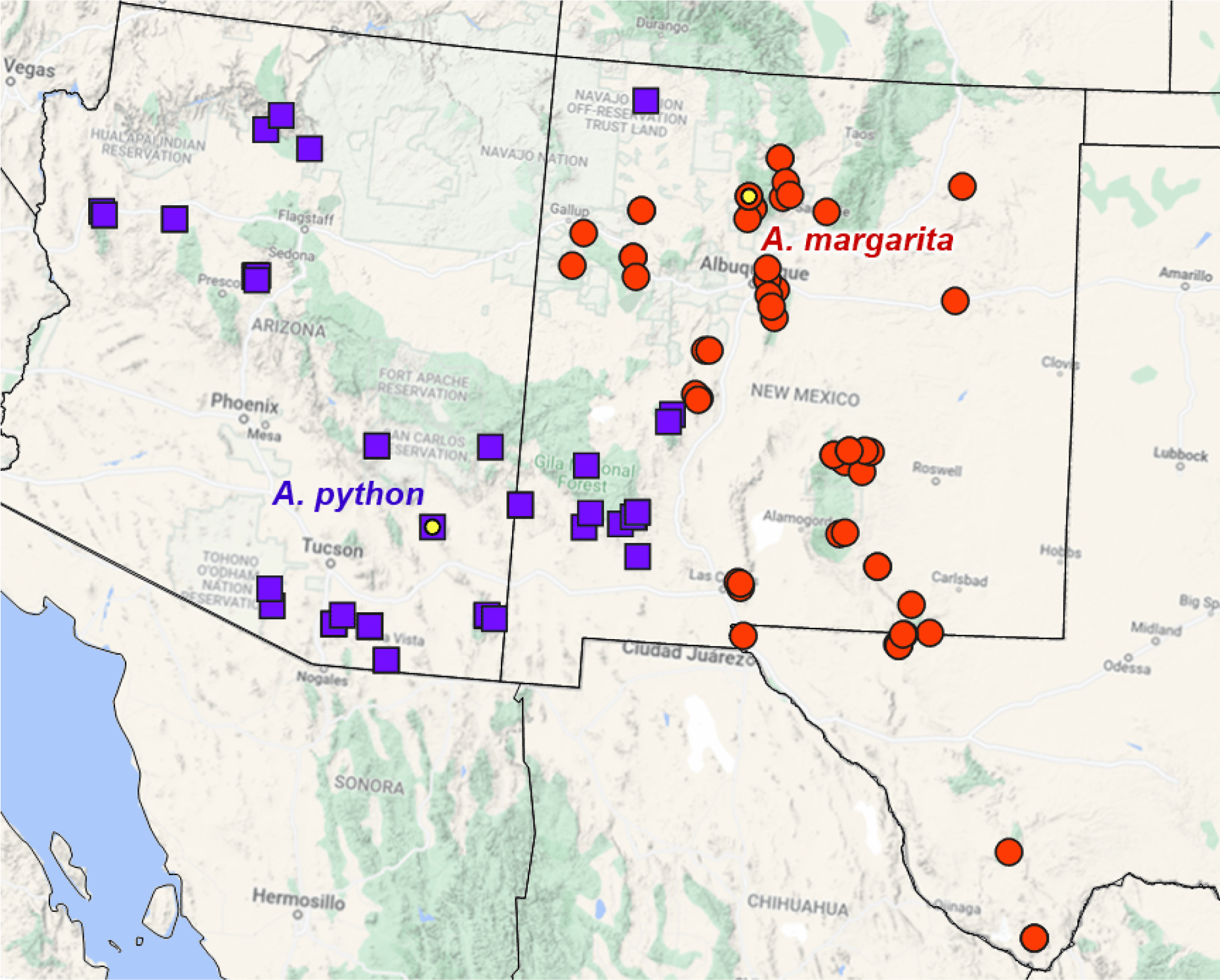
Occurrences of *A. python* (purple squares) and *A. margarita* (red circles) based on genomic analysis of >100 study specimens. Yellow dots = type localities. Latitude and longitude were estimated for most museum specimens.

To better understand how these similar skippers allocate habitats in Arizona and New Mexico, a closer look is helpful. The map in Figure 7 reveals that the very irregular Continental Divide almost completely separates these two skippers in the landscape. Python is fundamentally a creature of the western slope. It occurs below the Mogollon Rim in central Arizona and in sky islands of southeast Arizona. Its apparent northern limit lies within the Grand Canyon system of northern Arizona, extending upstream via tributaries into northwestern New Mexico. All these locations are west of the Divide and within the Colorado River watershed. It likely also occurs in Sonora, Mexico, but data are lacking. Uniquely, Python’s presence in southwest New Mexico is partly on the east slope of the Divide.

**Figure 7.**
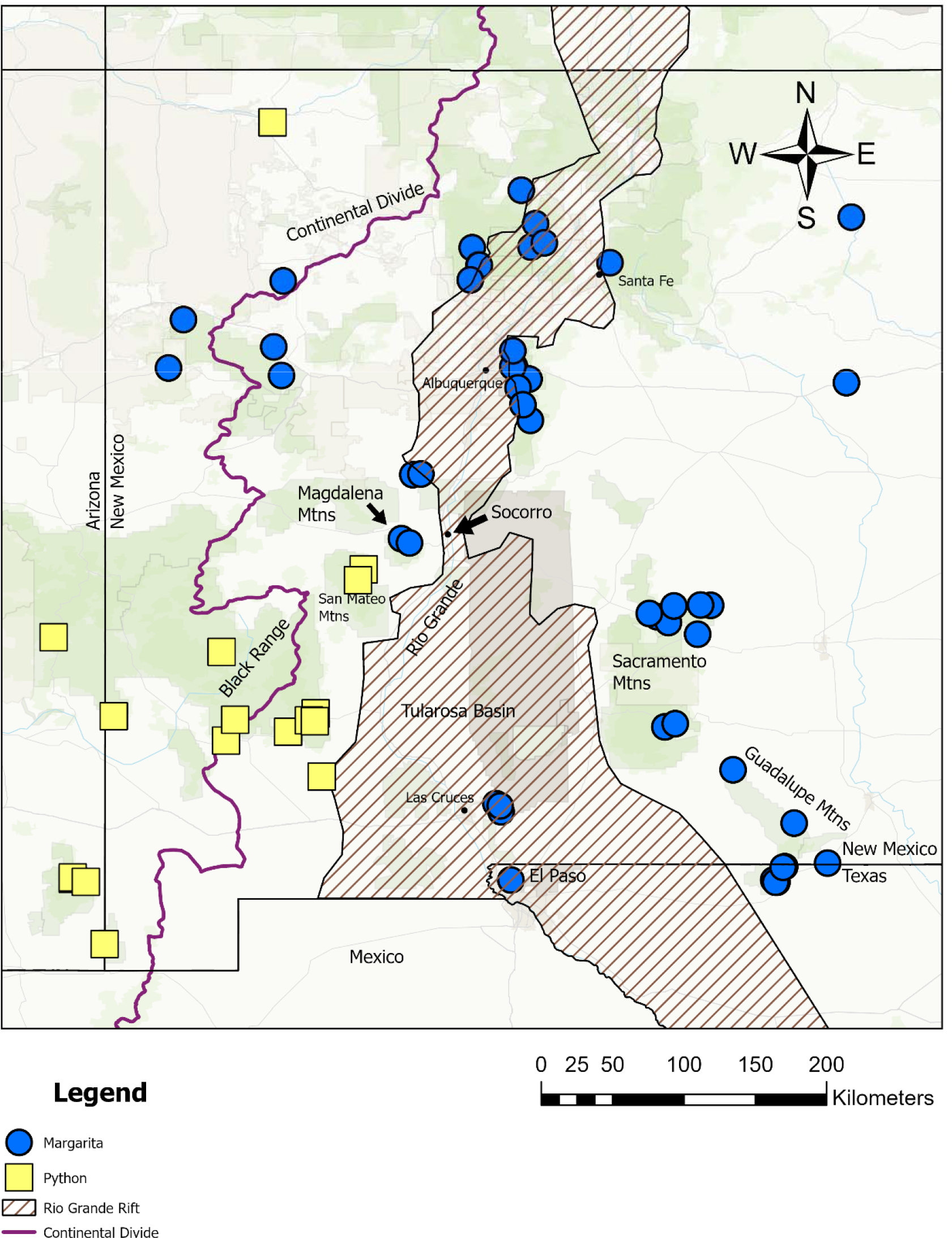
Study observations of Python and Margarita with reference to the Continental Divide and the Rio Grande Rift.

Margarita, in contrast, is fundamentally an east-slope organism. It inhabits much of northern, central and eastern New Mexico, extending into Trans-Pecos Texas. Its northern limit is in the Jemez and Sangre de Cristo Mountains. All those locations lie east of the Divide and, except for some internally drained basins, within the Rio Grande watershed, which ultimately drains to the Gulf of Mexico. Margarita has encroached westward across the Divide near Gallup, NM, and sparingly onto New Mexico’s eastern plains. Margarita undoubtedly occurs in adjacent Chihuahua, Mexico, but data are lacking.

Python and Margarita are mostly allopatric, but they approach each other in central New Mexico where the Magdalena Mountains and adjacent San Mateo Mountains each rise above 3050m (10,000ft) elevation. To date, analysis of multiple specimens from each range show that the San Mateos have only Python and the Magdalenas have only Margarita. The two species apparently function independently about 40km (25mi) from each other, separated only by Milligan Gulch. Evidence of introgression in one specimen from the Magdalenas hints at a common ancestor in the not-too-distant past.

### Paleobiogeography

Modern distributions of Python and Margarita (Fig. 6) resulted from their independent responses to Late Quaternary Period (Pleistocene and Holocene) climates and climate changes. The last full glacial climate persisted for about 100,000 years (Corrick et al. 2020). During those colder millennia, ice sheets spread across North America’s higher latitudes and most of what we now understand as mid-latitude plants and animals resided at more southerly latitudes or lower elevations. In the southwestern United States, abundant evidence for Pleistocene and Holocene ecosystems comes from radiocarbon-dated pollen, spores, phytoliths and microscopic charcoal from lake sediments, cave speleothems and fossilized packrat (*Neotoma* spp.) middens (Van Devender et al. 1984, Betancourt et al. 1990, 1993, 2001, Cordova and Johnson 2019; Reeves 1973; Hall and Valastro 1995; Arauza et al. 2016; Seersholm et al. 2020). Lepidoptera are poorly represented in regional Quaternary fossils (Elias 1992), but there are enough indisputable facts and reasonable suppositions to support a plausible explanation for how Python and Margarita arrived at their modern distributions.

During the last full glacial, Python and Margarita were reproductively isolated from each other. Glacial climates probably restricted Python to lower elevations on the west slope of the Divide, favoring southwest Arizona and northern Sonora, largely within the Colorado River watershed. Margarita would have been suppressed to lower sites in southeast New Mexico, Chihuahua, and west Texas, largely within the Rio Grande watershed. The series of down-dropped crustal blocks called the Rio Grande Rift is a narrow trough or trench in southern Colorado and northern New Mexico, but south of Socorro it widens to include the Jornada del Muerto and Tularosa Basins. The Tularosa Basin was the site of glacial Lake Otero and now hosts the largest gypsum dune field on Earth, likely a formidable barrier to east-west dispersal. The last full glacial was an opportunity for Python and Margarita to diverge genetically.

About 11,700 years ago, the rather abrupt change to warm, dry Holocene climates triggered northward and upslope range adjustments by many plants and animals. Major northward expansions for important woody plants including Ponderosa pine (*Pinus ponderosa*) and Piñon pine (*Pinus edulis*) are well documented (Norris et al. 2016, Cole et al. 2013), but postglacial dispersal of herbaceous plants and grasses, including hosts for *Atrytonopsis* larvae, still needs study (Avendaño-González et al. 2019). In regions of high physiographic relief like our study area, vertical range adjustments should have been frequent because many uplands offer complex habitats with opportunities to change elevation or slope aspect as needed (Noonan 1990).

Many southern uplands supporting Margarita or Python today likely did so during the last full glacial episode as well, though at lower elevations, and we suggest that modern occurrences of Python and Margarita are clues to their post-glacial dispersal routes (Fig. 7). For example, Margarita would have had a relatively unobstructed route north from Trans-Pecos Texas because the Guadalupe Mountains and Sacramento Mountains form a nearly continuous corridor of vertically complex habitats along the eastern margin of the 120km-wide (80mi-wide) Rio Grande Rift in that area. Once north of down-dropped crustal blocks forming the Tularosa Basin and Jornada del Muerto, Margarita had a clear path west to the Rio Grande. Near Socorro, narrowing of the Rift to about 32km (20mi) (Chapin and Seager 1975) allowed Margarita to cross the Rio Grande, leading to its present occupation of the Magdalena Mountains and setting the stage for its modern prevalence in northern New Mexico.

In contrast, Python’s postglacial upslope and northward expansion may have quickly encountered the insurmountable Mogollon massif. Eventually Python circumvented eastern buttresses of the Mogollon Mountains and dispersed east into southwest New Mexico through a broad gap in the Divide currently exploited by Interstate-10. When Python reached the Black Range it finally had a clear path north, but Margarita’s prior occupation of the Magdalena Mountains seems to have blocked further northward dispersal by Python in that area. Modern distributions of Margarita and Python, two distinct species, were thus established.

## ACKNOWLEDGMENTS

We thank institutional collection managers for their assistance in our efforts to locate, photograph, examine and sample specimens for further tests. Chuck Harp, at CSU’s C. P. Gillette Museum, is particularly appreciated for his ardent enthusiasm and helpfulness.

The authors are extremely grateful for the efforts of volunteers who worked hard to obtain supplemental specimens from additional locations than those represented in collections. Bill Dempwolf and Matthew Brown made multiple trips from neighboring states, drove many miles, burned much gasoline, and spent many hours in this effort. Conditions were poor in both years of their effort and the fact that they produced any specimens at all attests to their skills and perseverance. New Mexico butterflyers Gordon Berman, Simon Doneski, Bonnie Frey, Rebecca Gracey, Sajan KC, Renee Robillard, Marta Reece, Anisha Sapkota, Joe Schelling, Hira Walker, Natalie Wells and Rob Wu each contributed valuable time, effort and enthusiasm to the effort. Their curiosity, dedication and comradeship are much appreciated.

During more than 40 years of Lepidoptera studies in New Mexico, Richard Holland collected and accumulated more than 100 specimens of Python and Margarita (Toliver et al. 2001). In his field notes and on his specimen labels he characterized them as *Atrytonopsis python* pursuant to Miller and Brown (1981:47), a widely respected contemporary authority. Holland’s collection ultimately went to the Clyde P. Gillette Museum at Colorado State University at his request. Many specimens we examined and subjected to DNA analysis for our study were collected by Holland, whose positive influence on understanding New Mexico butterflies continues to be felt.

Mike Toliver provided thoughtful reviews of early drafts and flagged wing fringes as potentially helpful features.

## Data Availability Statement

Whole-genome shotgun datasets generated and analyzed in this study will be deposited in the NCBI database (https://www.ncbi.nlm.nih.gov) under BioProject PRJNAXXXXXXX

## Author Contribution Statement

SJC: Writing – original draft (lead); review and editing (equal); Conceptualization (lead); Investigation (lead); Analysis of Specimens; Data Curation. SMD: Writing – original draft (supporting); review and editing (equal); Visualization (equal); Investigation (supporting). JZ: Genomic Analysis (equal); Writing – original draft (supporting); review and editing (supporting). QC: Genomic Analysis (equal); Writing – original draft (supporting); review and editing (supporting). NVG: Writing – original draft (supporting); review and editing (equal); Visualization (equal); Genomic Analysis (equal); Resources.

## References

Arauza, H. M., A. R. Simms, L. C. Bement, B. J. Carter, T. Conley, A. Woldergauy, W. C. Johnson, & P. Jaiswal. 2016. Geomorphic and sedimentary responses of Bull Creek Valley (Southern High Plains USA) to Pleistocene and Holocene Environmental Changes. Quat. Res. 85(1): 118–132. DOI:10.1016/j.yqres.2015.11.006

Ashworth, A. C. 1973. Fossil Beetles from a Fossil Wood Rat Midden in Western Texas. Coleopterists Bulletin 27(3): 139–140.

Avendaño-González, M., J. F. Morales-Domínguez, & M. E. Siqueiros-Delgado. 2019. Genetic structure, phylogeography, and migration routes of *Bouteloua gracilis* (Kunth) Lag. ex Griffiths (Poaceae: Chloridoideae). Molecular Phylogenetics and Evolution 134: 50–60.

Bailowitz, R. & J. Brock. 2022. Southeastern Arizona Butterflies. Wheatmark Publ. Co. Tucson, AZ. 356 pp.

Betancourt, J. L., T. R. Van Devender, & P. S. Martin. 1990. Packrat Middens: The Last 40,000 Years of Biotic Change. University of Arzona Press. Tucson. DOI:10.2307/j.ctv21wj578

Betancourt, J.L., E. Pierson, & J. A. Fairchild-Parks. 1993. Influence of History and Climate on New Mexico Piñon-Juniper Woodlands. Pages 42–62 In Aldon, E. C. and D. W. Shaw (Tech. Coords.). Managing Piñon-Juniper Ecosystems for Sustainability and Social Needs. USDA For. Serv.. Gen Tech. Report RM-236.

Betancourt, J. L., K. A. Rylander, C. Peñalba, & J. L. McVickar. 2001. Late Quaternary vegetation history of Rough Canyon, south-central New Mexico, USA. Paleogeography, Paleoclimatology and Paleoecology 165(1-2): 71–95.

Burns, J. M. 2015. Speciation in an Insular Sand Dune Habitat: *Atrytonopsis* (Hesperiidae: Hesperiinae) – mainly from the Southwestern United States and Mexico – off the North Carolina Coast. Journal of the Lepidopterists’ Society 69(4): 275–292.

Cary, S. J., & M. E. Toliver. 2026. Butterflies of New Mexico. Available from Pajarito Environmental Education Center (https://peecnature.org/butterflies-of-new-mexico/). Accessed (September 9th, 2025)

Chapin, C. E. & W. R. Seager. 1975. Evolution of the Rio Grande rift in the Socorro and Las Cruces areas. New Mexico Geol. Soc. Guidebook 26: 297–321. DOI:10.56577/FFC-26.297 in: Las Cruces Country, Seager, W. R., R. E. Clemons, J. F. Callender, [eds.], New Mexico Geological Society 26th Annual Fall Field Conference Guidebook, 376 p. DOI:10.56577/FFC-26

Cole, K. L., J. F. Fisher, K. Ironside, & J. I. Mead. 2013. The biogeographic histories of *Pinus edulis* and *Pinus monophylla* over the last 50,000 years. Quaternary International 310(1):96–110.

Cordova, C. E. & W. C. Johnson. 2019. An 18ka to present pollen- and phytolith-based vegetation reconstruction from Hall’s Cave, south-central Texas, USA. Quat. Res. 92(2):497–518.

Corrick, E. C., R. N. Drysdale, J. C. Hellstrom, E. Capron, S. O. Rasmussen, X. Chang, D. Fleitmann, I. Couchoud, & E. Wolff. 2020. Synchronous timing of abrupt climate changes during the last glacial period. Science. 369 (6506): 963–969. DOI:10.1126/science.aay5538.

Edwards, W. H. 1882. Descriptions of new species of diurnal Lepidoptera taken by Mr. H. K. Morrison, at Fort Grant and in Graham Mountains, Arizona, 1882. Papilio 2(8): 136–143. —1884. Descriptions of new species of butterflies from Arizona. Papilio 4(3):53–58.

Elias, S. A. 1987. Paleoenvironmental Significance pf Late Quaternary Insect Fossils from Packrat Middens in South-Central New Mexico. Southwest Naturalist 32(3):383–390.

Elias, S. A. 1992. Late Quaternary zoogeography of the Chihuahuan Desert insect fauna, based on fossil records from packrat middens. Journal of Biogeography 19(3): 285–297. DOI:10.2307/2845452

Ficarrotta, V., J.J. Hanly, L.S. Loh, C.M. Francescutti, A. Ren, K. Tunström, C.W. Wheat, A.H. Porter, B.A. Counterman, & A. Martin. 2022. A genetic switch for male UV iridescence in an incipient species pair of sulphur butterflies, Proc. Natl. Acad. Sci. U.S.A. 119 (3) e2109255118, DOI:10.1073/pnas.2109255118.

Godman, F. D. & O. Salvin. 1900. Biologia Centrali Americana. Zoología, Insecta, Lepidoptera Rhopalocera. Vols. II,III. Dulau & Co., Bernard Quaritch. London. Pp.

Hall, S. & S. Valastro. 1995. Grassland vegetation in the southern great plains during the last glacial maximum. Quat. Res. 44(2): 237–245. DOI:10.1006/QRES.1995.1068

Hoang, D. T., O. Chernomor, A. von Haeseler, B. Q. Minh, & L. S. Vinh. 2018. UFBoot2: improving the ultrafast bootstrap approximation. Molecular Biology and Evolution 35(2): 518–522.

Li, W., Q. Cong, J. Shen, J. Zhang, W. Hallwachs, D. H. Janzen, & N. V. Grishin. 2019. Genomes of skipper butterflies reveal extensive convergence of wing patterns. Proceedings of the National Academy of Sciences of the United States of America 116(13): 6232–6237. DOI: 10.1073/pnas.1821304116

McNally, P. 2020. Butterflies of the Central Arizona Highlands. ECO Publishing. Rodeo, NM. 299 pp.

Mielke, O. H. H. 2005. Catalogue of the American Hesperioidea: Hesperiidae (Lepidoptera), Vol. 4, Hesperiinae 1: Adlerodea Lychnuchus. Curitiba: Soc. Brasileria de Zoologia.

Miller, L. D. & F. M. Brown. 1981. A Catalog/Checklist of the Butterflies of America, north of Mexico.

Nguyen, L. T., H. A. Schmidt, A. von Haeseler, & B. Q. Minh. 2015. IQ-TREE: a fast and effective stochastic algorithm for estimating maximum-likelihood phylogenies. Molecular Biology and Evolution 32(1): 268–274.

Noonan, G.R. 1990. Biogeographical patterns of North American *Harpalus* Latreille (Insecta: Coleoptera: Carabidae). J. Biogeogr. 17: 583–614.

Norris, J. R., J. Betancourt, L. Julio, & S. T. Jackson. 2016. Late Holocene Expansion of Ponderosa Pine (*Pinus ponderosa*) in the Central Rocky Mountains. J. Biogeogr. 43: 778–790.

Rambaut, A. 2018. FigTree, version 1.4.4. Available at http://tree.bio.ed.ac.uk/software/figtree/ [Accessed May 2026].

Reeves, C.C. Jr. 1973. The Full Glacial Climate of the Southern High Plains, Texas. Jour. Geol. 84(6):693–704.

Seersholm, F. V., D. J. Werndly, A. Grealy, T. Johnson, E. M. Early, E. L. Lundelius, B. Winsborough, G. E. Farr, R. Toomey, A. J. Hansen, B. Shapiro, M. R. Waters, G. McDonald, A. Linderholm, T. W. Stafford & M. Bunce. 2020. Rapid range shifts and megafaunal extinctions associated with late Pleistocene climate change. Nat Commun 11, 2770. DOI:10.1038/s41467-020-16502-3.

Skinner, H. A. New *Pamphila* from New Mexico (Lepidoptera). Can. Entomol. 45:426–427.

Toliver, M. E., R. Holland, & S. J. Cary. 2001. Distribution of Butterflies in New Mexico. 3^rd^ Ed. Published by the authors. 449 pp. + appendices.

Van Devender, T. R., J. L. Betancourt, & M. Wimberly. 1984. Biogeographic Implications of a Packrat Midden Sequence from the Sacramento Mountains, South-central New Mexico. Quaternary Research 22(3): 344–360.

Wilkins, D.E. & D. R. Currey. 1997. Timing and Extent of Late Quaternary Paleolakes in the Trans-Pecos Closed Basin, West Texas and South-Central New Mexico. Quat. Res. 47(3): 306–315. DOI:10.1006/qres.1997.1896

